# A Facilitated Diffusion Mechanism Establishes the Drosophila Dorsal Gradient

**DOI:** 10.1101/057091

**Authors:** Sophia N. Carrell, Michael D. O’Connell, Amy E. Allen, Stephanie M. Smith, Gregory T. Reeves

## Abstract

The transcription factor NF-κB plays an important role in the immune system as an apoptotic and inflammatory factor. In the *Drosophila melanogaster* embryo, a homolog of NF-ΚB called Dorsal (dl) patterns the dorsal-ventral (DV) axis in a concentration-dependent manner. During early development, dl is sequestered outside the nucleus by Cactus (Cact), homologous to IκB. Toll signaling at the ventral midline breaks the dl/Cact complex, allowing dl to enter the nucleus where it transcribes target genes. Here we show that dl accumulates on the ventral side of the embryo over the last 5 cleavage cycles and that this accumulation is the result of facilitated diffusion of dl/Cact complex. We speculate that the predominant role for Cact in DV axis specification is to shuttle dl towards the ventral midline. Given that this mechanism has been found in other, independent systems, we suggest it may be more prevalent than previously thought.

## Introduction

In a developing organism, tissues are patterned by long-range signaling enacted through morphogen concentration gradients that carry the positional information necessary to control gene expression in a spatially-dependent fashion. In the *Drosophila* blastoderm, the transcription factor Dorsal (dl) acts as a morphogen to regulate the spatial patterning of greater than 50 genes along the dorsal-ventral (DV) axis (Moussian and Roth, 2005; Reeves and Stathopoulos, 2009). However, despite its centrality to development, the mechanism that controls the formation of the dl nuclear concentration gradient at the whole-embryo scale is not well understood.

At the single nucleus/cell level, dl is sequestered to the cytoplasm in an inactive complex with the IκB homolog Cactus (Cact). On the ventral side of the embryo, Toll signaling results in the degradation of Cact, freeing dl to enter the nucleus where it regulates transcription. This mechanism, combined with a ventral-to-dorsal gradient of Toll signaling, would seem sufficient to develop a gradient of nuclear dl concentration (Roth et al., 1989). However, taken alone, this mechanism would result in a local depletion of dl from the cytoplasm surrounding the ventral nuclei to create a counter-gradient in cytoplasmic dl, which is in contrast to the observation that dl accumulates both in the nuclei and in the cytoplasm on the ventral side of the embryo over time (Reeves et al., 2012) (see Figure 1A–D).

**Figure 1:**
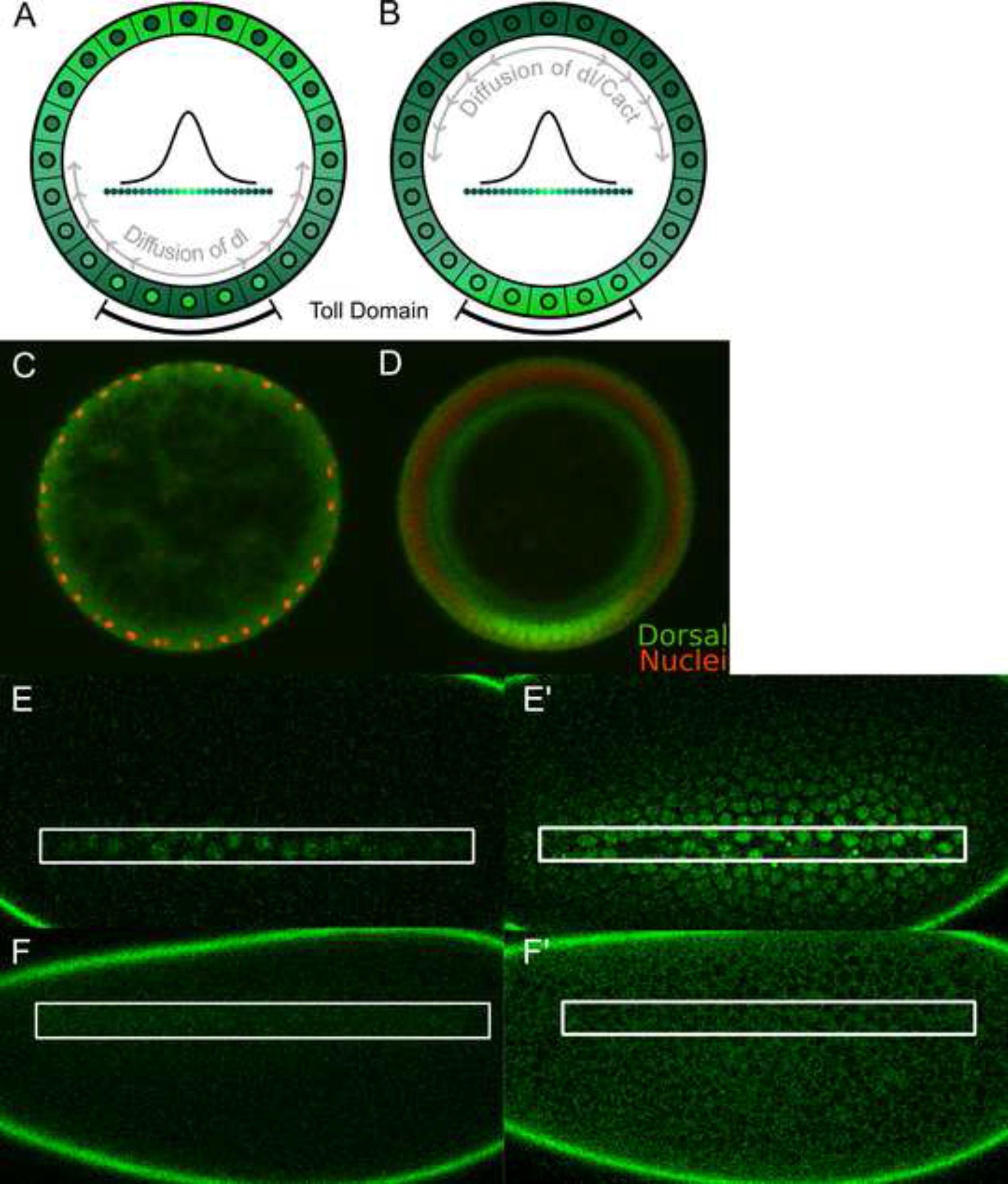
Dorsal Accumulates on the Ventral Side of the Embryo. (A) Current research suggests that dl does not rapidly diffuse from nucleus to nucleus and therefore it is locally depleted from the area around each nucleus. (B) We propose that total dl accumulates at the ventral midline due to diffusion. “Toll Domain” indicates that active Toll receptors are being produced on the ventral side of the embryo. (C) and (D) Cross-sections of wild type embryos, stained for dl and nuclei. (C) dl is equally distributed throughout when the embryo is young. (D) Total dl has accumulated at the ventral midline by late nc 14. Compare to (B). (E) and (F) dl-paGFP results indicate that dl diffuses throughout the embryo. Activation area in the white box. Anterior to the left. (E and E′) Activation near the ventral midline approx. (E) 3 minutes and (E′) 90 minutes after first activation. (F and F′) Activation near the dorsal midline approx. (F) minutes and (F′) 90 minutes after first activation. See also Movies S1 and S2.

In this study, we investigate the mechanism of dl nuclear concentration gradient formation at the whole-embryo scale. We use a computational model to guide the design and interpret the outcome of experiments that distinguish between competing hypotheses. Our experimental and computational datasupport the hypothesis that a mechanism of facilitated diffusion is responsible for the global formation of the dl nuclear concentration gradient. Under this “shuttling” hypothesis, a dorsal-to-ventral concentration gradient of cytoplasmic dl/Cact complex develops as the result of Toll-mediated degradation of Cact, which drives the overall flux of dl towards the ventral side of the embryo. Besides ventrally-directed dl accumulation, our computational model predicts that the shuttling mechanism can account for several other previously inadequately explained phenotypes, such as duplicated dl gradient peaks, regions of depleted dl, dl heterozygous phenotypes, and widened dl-GFP gradients (Huang et al., 1997; Liberman et al., 2009; Reeves et al., 2012; Roth and Schüpbach, 1994). We have experimentally verified these predictions that a facilitated diffusion mechanism is responsible for the formation of the dl nuclear concentration gradient in the developing *Drosophila* embryo.

The mechanism of facilitated diffusion has been previously described and is responsible for gradient formation in other systems, such as the Dpp/BMP signaling pathway (Ben-zvi et al., 2008; Eldar et al., 2002; Marqués et al., 1997; Mizutani et al., 2005; Shimmi et al., 2005) and has also been suggested for the formation of the Spätzle gradient upstream of Toll signaling (Haskel-Ittah et al., 2012). The dl/Cact system has each of the features required for facilitated diffusion (Shilo et al., 2013): (1) the primary molecule (dl) binds to a “carrier” molecule (Cact) that protects it from capture, (2) the primary/carrier complex is diffusible (due to the syncytial state of the early embryo), and (3) the complex is broken in a spatially-dependent manner (through Toll signaling on the ventral side of the embryo). If these features are in place, the primary molecule (dl) accumulates over time at the site of complex degradation (the ventral side of the embryo). Therefore, we conclude that Cact performs a role in dl gradient formation additional to regulating dl’s entry into the nucleus: shuttling dl to the ventral side to form the mature gradient.

## Results

### Dorsal accumulation on the ventral side of the embryo results from movement of the dl/Cact complex

Initially, Dorsal is uniformly distributed along the DV axis of the developing embryo (Figure 1C); during nuclear cycles 11-14, it accumulates on the ventral side (Figure 1B,D). This observation is consistent with previously-published fluorescent images of anti-dl immunostainings in fixed embryos and in images of dl-GFP fluorescence in live embryos (Kanodia et al., 2009; Liberman et al., 2009; Reeves and Stathopoulos, 2009; Reeves et al., 2012). The mechanism for this overall polarization of total dl in the embryo could be asymmetric degradation or production; however, our modeling work and experiments favor an overall flux of dl to the ventral side of the embryo. Using a photoactivatable GFP (paGFP) tag (Patterson and Lippincott-Schwartz, 2002), we noticed that dl appears in regions of the embryo over 8-10 nuclei away from the site of paGFP activation in the time span of 90 minutes. Given a cell diameter (i.e., the average distance between the centroids of two neighboring nuclei) of roughly 7 μm, this translates to an effective diffusivity of ∼0.5-0.9 μm^2^/s and a time scale to cross one cell diameter that is on the order of minutes. When activated near the ventral midline, dl-paGFP diffuses from its location of activation and fills adjacent nuclei (Figure 1E, Movie S1). When activated on the dorsal side of the embryo, dl-paGFP diffuses and shows a typical pattern of exclusion from nuclei in more dorsal regions and moderate uptake into the nuclei in more lateral regions (Figure 1F, Movie S2). As discussed previously, this flux of dl could be due to diffusion of free dl, dl/Cact complex, or both species (O’Connell and Reeves, 2015).

To deduce which process(es) may be responsible for the overall flux of dl to the ventral side of the embryo, we used a model of dl/Cact interactions (Figure 2A,B; see also Supplementary Experimental Procedures) and examined the equation for total dl (see Experimental Procedures). From this equation, we see that only two species may be contributing to the overall movement of dl to the ventral side of the embryo: free cytoplasmic dl and Cact-bound cytoplasmic dl. Note that this is a generic prediction of our model equations and does not rely on any specific model parameter values.

**Figure 2:**
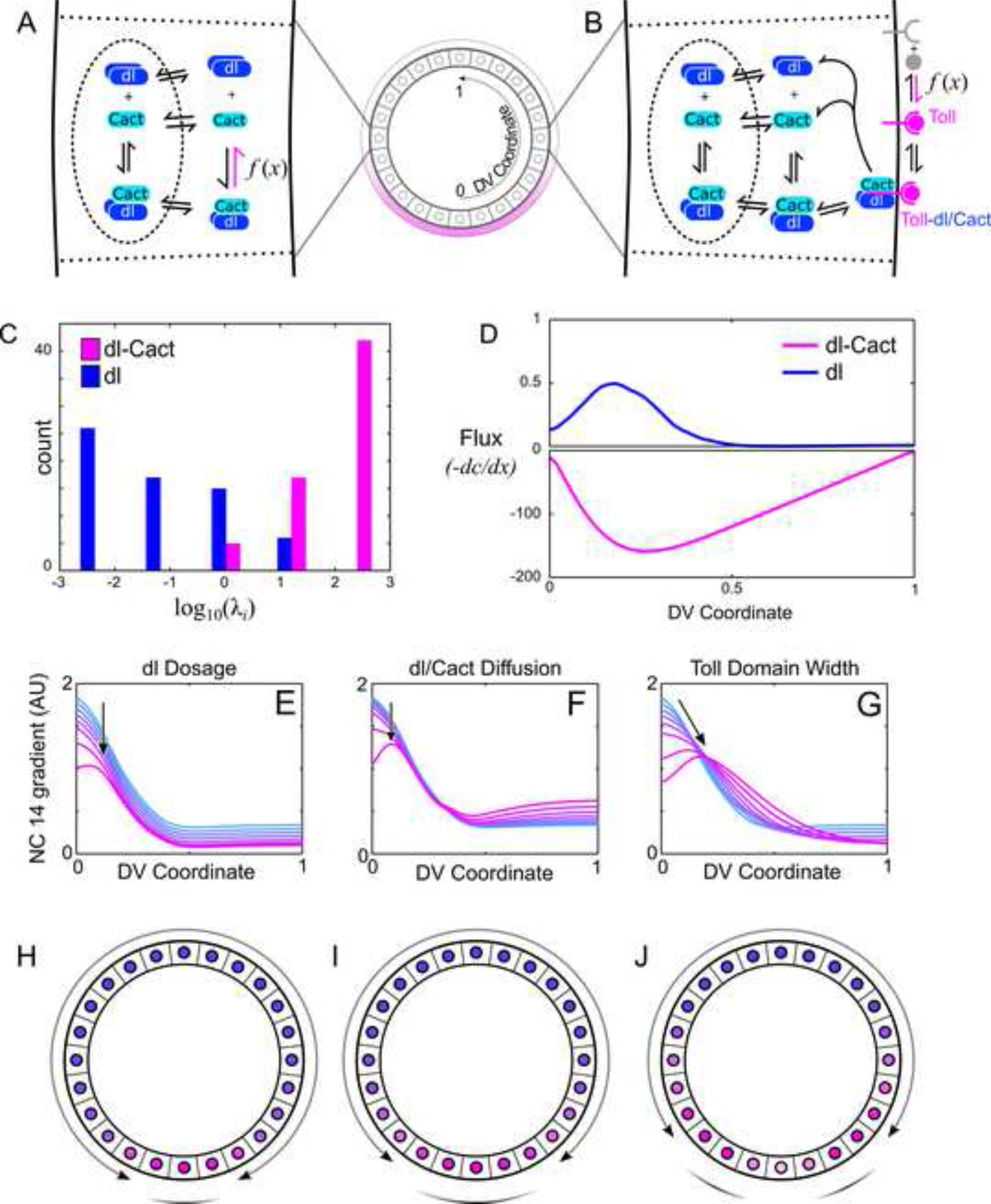
The Model Predicts a Shuttling Mechanism and Mutant Phenotypes. (A) The previously published model of dl gradient dynamics in which the active Toll region was modeled phenomenologically (catalytic activity labeled *f(x)*). (B) Expanded model used in the current study. Active Toll receptors are explicitly accounted for (produced at a rate *f(x)*). (C) Evolutionary optimization favors high values for dl/Cact diffusion (dimensionless parameter *λ_dl/Cart_*) and low values for free dl diffusion (*λ_dl_*). (D) The model predicts dl/Cact diffusion toward the ventral midline (negative values) is far greater than free dl diffusion away from the ventral midline (positive values). (E-G) Perturbing a set of representative parameters to reduce rate of shuttling results in a hallmark progression of phenotypes. The dl gradient first widens (width parameter o increases), then flattens at the peak, then obtains a split peak by decreasing the dosage of dl (E), increasing dl/Cact diffusion (F), and increasing the width of the active Toll domain (G). Arrows indicate direction of change upon decreasing the rate of shuttling. (H-J) Illustrations of the hallmark progression of phenotypes associated with shuttling: the peak expands (H), then flattens (I), then splits into two peaks (J). Arrows indicate the direction of flux of total dl.

### Model analysis and experimental predictions

To determine which species, free dl or dl/Cact complex, is responsible for the accumulation of dl on the ventral side of the embryo, we analyzed the full set of model equations (Equations 1-8 in Supplemental Experimental Procedures) and made the model consistent with spatiotemporal data of dl gradient dynamics (Reeves et al., 2012) using evolutionary optimization. Analysis of the resulting parameter sets reveals that, to be consistent with the data, the rate of dl/Cact complex diffusion is generally much faster compared to that of free dl (Figure 2C).

We then analyzed the direction in which dl and dl/Cact complex diffuse in the model. According to Fick’s law, diffusive flux (typically denoted *J*) refers to the movement of a solute from areas of high concentration to areas of low concentration at a rate proportional to its concentration gradient, *dC/dx*, yielding *J ∝ dC / dx*. The negative sign accounts for the fact that diffusive flux is in the direction of decreasing concentration. By convention, we define our x-axis to be equal to zero at the ventral midline and one at the dorsal midline (see Figure 2A,B); thus, a negative value for the flux indicates movement toward the ventral midline, and a positive value indicates movement toward the dorsal midline.

Our modeling results indicate that dl/Cact complex preferentially moves towards the ventral midline, while free dl moves away (Figure 2D). Therefore, the ventrally-directed flux of total dl must be due to diffusion of dl/Cact complex, which results from Toll signaling acting as a sink at the ventral midline (see also Figure 1A,B). In this way, Cact shuttles dl against its concentration gradient, thereby allowing an accumulation of nuclear dl at the ventral midline where it is required for proper formation of the dl gradient.

The model used here is an extension of a previously published model (O’Connell and Reeves, 2015), in which the catalytic activity of Toll signaling was modeled phenomenologically (Figure 2A,B). We updated the model to explicitly account for both activated Toll receptors and Toll receptors bound to dl/Cact complex (Equations 7 & 8, respectively) to allow for the possibility that Toll activity is limiting. This possibility was precluded in the previous model, because the rate of Toll-mediated Cactus degradation was proportional to the concentration of dl/Cact, which created a potentially unlimited sink.

Further analysis of our model showed that the mechanism of facilitated diffusion can be tested by altering three experimentally-tunable biophysical parameters to slow the mechanism of facilitated diffusion: (1) decreasing the dl dosage, (2) lowering the diffusivity of dl or Cact, and (3) increasing the width of the active Toll domain (Figure 2E–G). The model predicts that these perturbations will each result in a hallmark progression of phenotypes (depending on the strength of the perturbation): the dl gradient first widens, then becomes flat at the top, then splits into two peaks (Figure 2H–J).

Furthermore, combinations of these perturbations have synergistic effects. We conducted experiments to address all three parameters, and found that they supported our hypothesis.

### Decreasing dl dosage widens and flattens the dl gradient

In embryos from mothers heterozygous null for dl (hereafter referred to as *dl/+* embryos), the shape of the gradient becomes slightly wider, flatter, and lower in amplitude, with the slope maintained in the lateral regions but peak levels of the gradient flattened (Liberman et al., 2009) (Figure 3). From a functionality perspective, this phenotype is striking because the portion of the dl gradient that is lost (the peak concentration) is superfluous as any level above the threshold level for high-concentration target genes will result in expression, while the portion of the gradient that is necessary for patterning gene expression boundaries—the graded portion between 20% and 45% DV position—is maintained.

**Figure 3:**
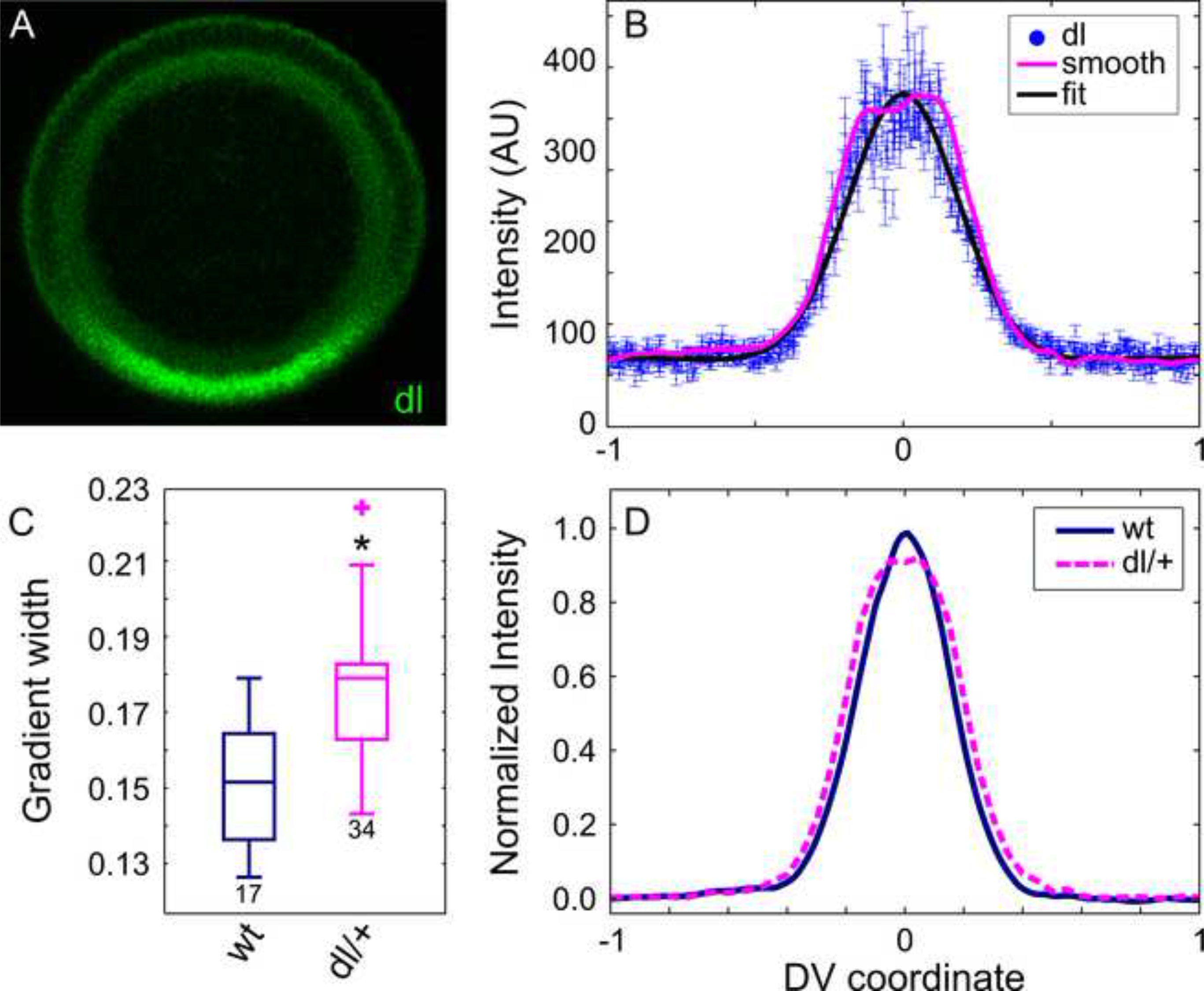
Embryos from Mothers Heterozygous for dl Have a Different-Shaped dl Gradient. (A) Crosssection of a NC 14 embryo from a mother heterozygous for *dl* stained for dl. (B) Plotted nuclear intensity of embryo in (A) as a function of DV coordinate (each blue dot is one nucleus). Error bars indicate SEM of nuclear intensity. Pink curve is the smoothed data; black curve is a fit of the data to a Gaussian. (C) Box plot of gradient widths (σ) for embryos from mothers with 2 (wt) and 1 (dl/+) copies of dl. Here and elsewhere in the paper, boxes indicate interquartile range (IQR): from the 25^th^ to the 75^th^ percentile of the data, whiskers extend a maximum of 1.5 times the width of the IQR from the box, and outliers (plus signs) are defined as points that lie outside the whiskers. Numbers indicate sample size. Asterisk indicates statistical significance of p<10^−4^. (D) Normalized average plot of dl gradient in embryos from (C). See also Figure S1.

To explain this effect, one simple hypothesis is that the ventral-most nuclei *would have* reached normal dl levels, but only if there were enough dl present. Since the total amount of dl is compromised, the ventral most nuclei reach a lower-than-normal concentration of dl. However, models of dl gradient formation that lack the shuttling mechanism predict the loss of 50% dl nuclear concentration globally in dl/+ embryos, rather than a spatially patterned loss (Figure S1).

In contrast, a model based on the shuttling mechanism makes sense of this phenotype. In wild type embryos, the smooth peak of the dl gradient is obtained due to a saturation effect. There is enough dl/Cact complex to saturate active Toll receptors in the ventral-lateral regions of the embryo, which allows a significant flux of dl/Cact complex to reach the ventral midline and results in a smoothly- peaked, Gaussian-like curve. In embryos with half the normal dl dosage, active Toll receptors in the tails of the gradient are not saturated; therefore, less dl/Cact complex arrives at the ventral midline, and these embryos never accumulate a smooth peak of the dl gradient (Figure 3B,D).

### Decreased diffusion of dl/Cact complex widens the dl gradient

Other researchers have generated several GFP-tagged versions of dl in order to study its dynamics in living embryos (Delotto et al., 2007; Reeves et al., 2012). In each case, tagging dl with an extra protein moiety caused the dl gradient to expand. If the gradient develops via a diffusion mechanism, this result would seem counterintuitive. One would expect that increasing the size of a molecule would decrease its diffusivity, and thus restrict the spatial range of the gradient. On the other hand, a shuttling mechanism would predict the opposite: that lowering the diffusivity of dl/Cact complex would widen the gradient. To test this prediction, we analyzed dl and Cact constructs tagged with proteins that either dimerize or tetramerize in order to form large clusters of dl/Cact complex.

Other studies have shown that tagging dl with a monomeric Venus (dl-mVenus) causes the dl gradient to widen (width parameter σ = 0.16 ± 0.01; compare to σ = 0.14 ± 0.01 for wild type),while a GFP tag causes a much greater widening (σ = 0.20 ± 0.02) (Liberman et al., 2009; Reeves et al., 2012). We surmised that the difference between these scenarios is that Venus is an obligate monomer while the GFP construct weakly dimerizes (Zacharias et al., 2002). To carefully measure the effect of dimerization of the protein tag on the width of the gradient, we repeated these experiments in order to control for differences in fluorescent protein tag, protocol, researcher, and equipment so as to have a direct comparison for our work in new fly lines. First, we analyzed the gradient in embryos that had one endogenous copy of dl replaced with dl tagged with the weakly-dimerizing GFP (dl-dGFP) (Reeves et al., 2012). We found that the dl gradient in these embryos widened significantly (σ = 0.21 ± 0.03) as compared to wild type (Figure 4A,B).

**Figure 4:**
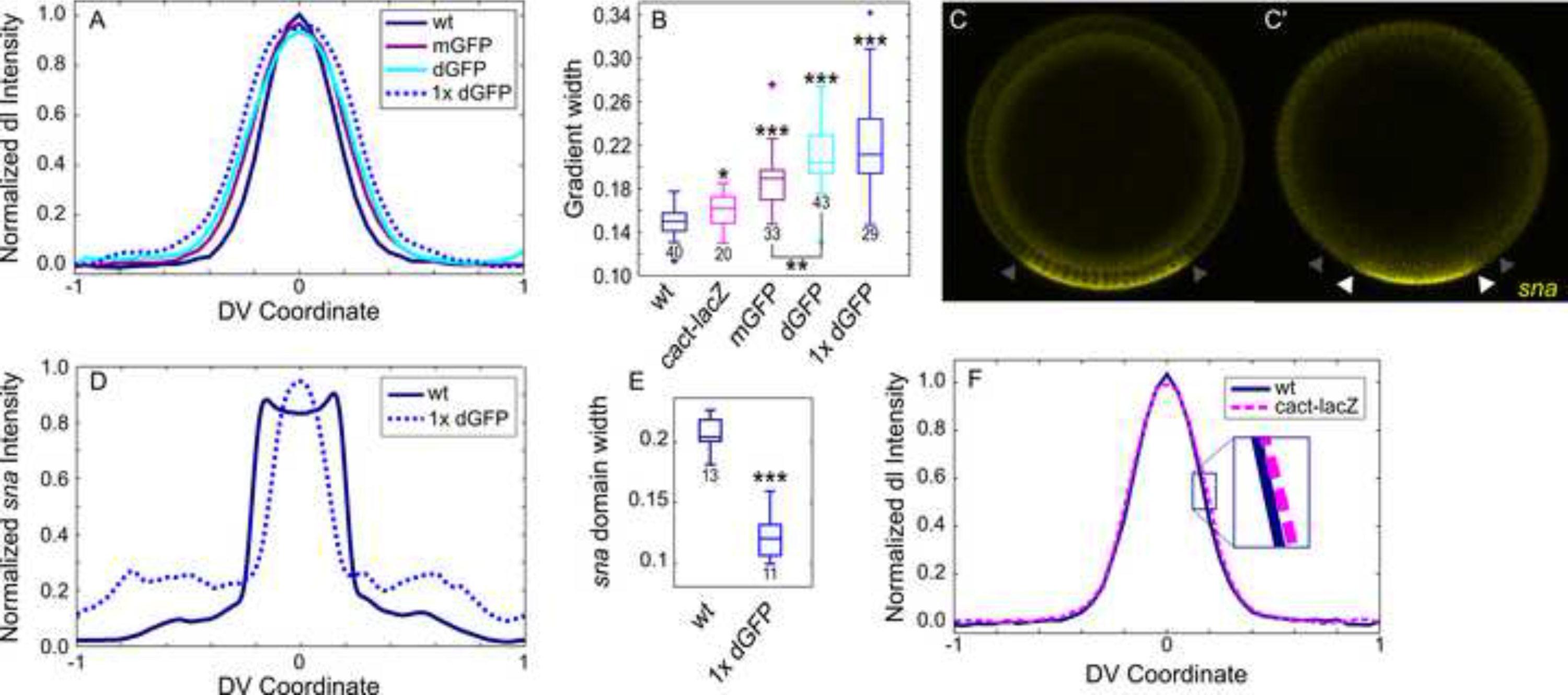
Decreasing Diffusion of dl/Cact Increases the Width of the dl Gradient. (A) Normalized average plots of the dl gradient in embryos with 2 copies of wt dl (wt), 1 copy of wt dl and 1 copy of dl-mGFP (mGFP), 1 copy of wt dl and 1 copy of dl-dGFP (dGFP), and 1 copy of dl-dGFP (1× dGFP). (B) Box plot of gradient widths (σ) for embryos shown in (A) and (F). (C) and (C') *sna* domain in wt (C) and 1x dGFP (C′) embryos. Gray arrowheads indicate the wt domain of *sna;* white indicate the *sna* domain in 1× dGFP embryo. (D) Normalized average plots of the *sna* intensity in wt and 1× dGFP embryos. (E) Box plot of *sna* domain width for embryos shown in D. (F) Normalized average plots of the dl gradient in embryos from mothers with 2 copies of wt *cact* (wt) and 1 copy of wt *cact* and 1 copy of *cact-lacZ* (cact-lacZ). Inset shows more clearly that cact-lacZ is slightly wider than wt. Asterisks indicate statistical significance (*p<0.02; **p<10^−3^; ***p<10^−11^) from wt unless otherwise noted. Numbers indicate sample size. See also Figure S2.

The weak dimerization of GFP can be abolished by the A206K mutation (Zacharias et al., 2002); therefore, we designed dl-GFP constructs with the obligate monomer version of GFP (dl-mGFP) in order to investigate the effect of increasing the mass of dl without causing dimerization. Embryos carrying these constructs have gradients slightly wider than wild type (σ = 0.19 ± 0.03), comparable to our measurements in embryos with the monomeric Venus tag (σ = 0.18 ± 0.03) (Figure 4B, see also Figure S2A,B). We expect that the widening effect in embryos that carry a monomeric tag is likely due to the increased mass of the dl protein and not the dimerization of the tag. We also designed dl-Venus constructs that had weak dimerization restored due to a K206A mutation (dl-dVenus); embryos carrying this dl-dVenus transgene had dl gradients that were significantly wider than those in embryos carrying the dl-mVenus transgene (σ = 0.19 ± 0.02), matching the trend seen in dl-mGFP and dl-dGFP embryos (see Figure S2 A,B). Taken together, these data indicate that the weak dimerization of the fluorescent tags, which presumably slows diffusion of dl/Cact complex, is the key factor in widening the gradient.

It was also observed that a single copy of dl-dGFP was unable to complement null mutations in the endogenous dl gene; two copies are necessary to rescue the mutant (Liberman et al., 2009; Reeves et al., 2012). This result contrasts dl-mVenus, which complements at one copy. Our shuttling hypothesis explains this phenotype, as embryos from *dl* mothers carrying one copy of dl-dGFP have two defects in shuttling: a lower copy number of dl and a reduced diffusivity of the dl/Cact complex. These embryos have highly widened gradients (σ = 0.22 ± 0.04) (see Figure 4A,B and Figure S2C,D). We surmised that these gradients are perturbed enough to severely disrupt gene expression, perhaps not becoming strongly-peaked enough to turn on high-threshold genes, such as *snail (sna)*. Therefore, we analyzed *sna* expression in embryos from *dl; dl-dGFP/+* mothers, and found that these embryos either lacked *sna* expression or expressed *sna* in a very narrow domain (Figure 4C–E, see also Figure S2D), confirming our hypothesis that the double perturbation results in a breakdown of the typically robust patterning system.

To investigate the effect of larger complexes on dl/Cact diffusion, we evaluated the effect of a *dl-lacZ* transgene on the dl gradient (Govind et al., 1992). As the protein product of *lacZ,* β-galactosidase (β-gal), tetramerizes, we expect that this fusion will slow the diffusion of dl to a greater extent than dGFP. However, we found that two copies of this transgene in a *dl/+* background were needed in order to see an effect on the width of the dl gradient, which widened as predicted by the shuttling model (See Figure S2B,E). This is likely due to the fact that *dl-lacZ* acts as an antimorphic allele (Govind et al., 1992); such dl-βgal fusion proteins are suspected to be expressed at very low levels in surviving fly lines (Govind et al., 1996).

In order to study the effect of decreased Cact diffusion on dl gradient formation, we examined embryos from mothers carrying a single copy of a *cact-lacZ* transgene in a heterozygous *cact* background (Fernandez et al., 2001). We expect that this fusion will slow diffusion of the dl/Cact complex without affecting the diffusion of free dl. In these embryos, the dl gradient is expanded (Figure 4B,F), indicating an inability of Cact to properly shuttle dl to the ventral midline. In a naïve model, where Cact does not shuttle dl and there is no flux toward the ventral midline, we do not expect changing the diffusion of Cact to have such an effect on the dl gradient. It is important to note that, under the typical morphogen gradient phenomenon, decreasing the diffusivity of the morphogen would result in a narrower (more concentrated) gradient. Since our results consistently show that the dl gradient widens rather than narrows when the diffusivity of dl or Cact is reduced, we conclude that dl/Cact complex, and not free dl, is the dominant diffusive species.

### Increasing the width of the active Toll domain results in a split peak of dl

The shuttling hypothesis predicts that widening the active Toll domain results first in a widened, then flattened, then split dl gradient (Figure 2H–J). As the extent of the Toll domain is controlled by Gurken/EGFR signaling during oogenesis (Schüpbach, 1987; Sen et al., 1998), we analyzed embryos from mothers carrying a hypomorphic EGFR allele (*egfr^t1^*) (Roth and Schüpbach, 1994). We found that embryos from mothers heterozygous for this allele have significantly widened dl gradients, and most (10/12) embryos from homozygous mothers have gradients so wide that a split peak forms (see Figure 5). This result is consistent with previous reports that various *gurken* and *egfr* mutations generate a duplicated dl gradient as measured by dl staining, Twist staining, and sites of ventral furrow formation (Roth and Schüpbach, 1994), which is not readily explained in the absence of a shuttling phenomenon (Meinhardt, 2004; Moussian and Roth, 2005).

**Figure 5:**
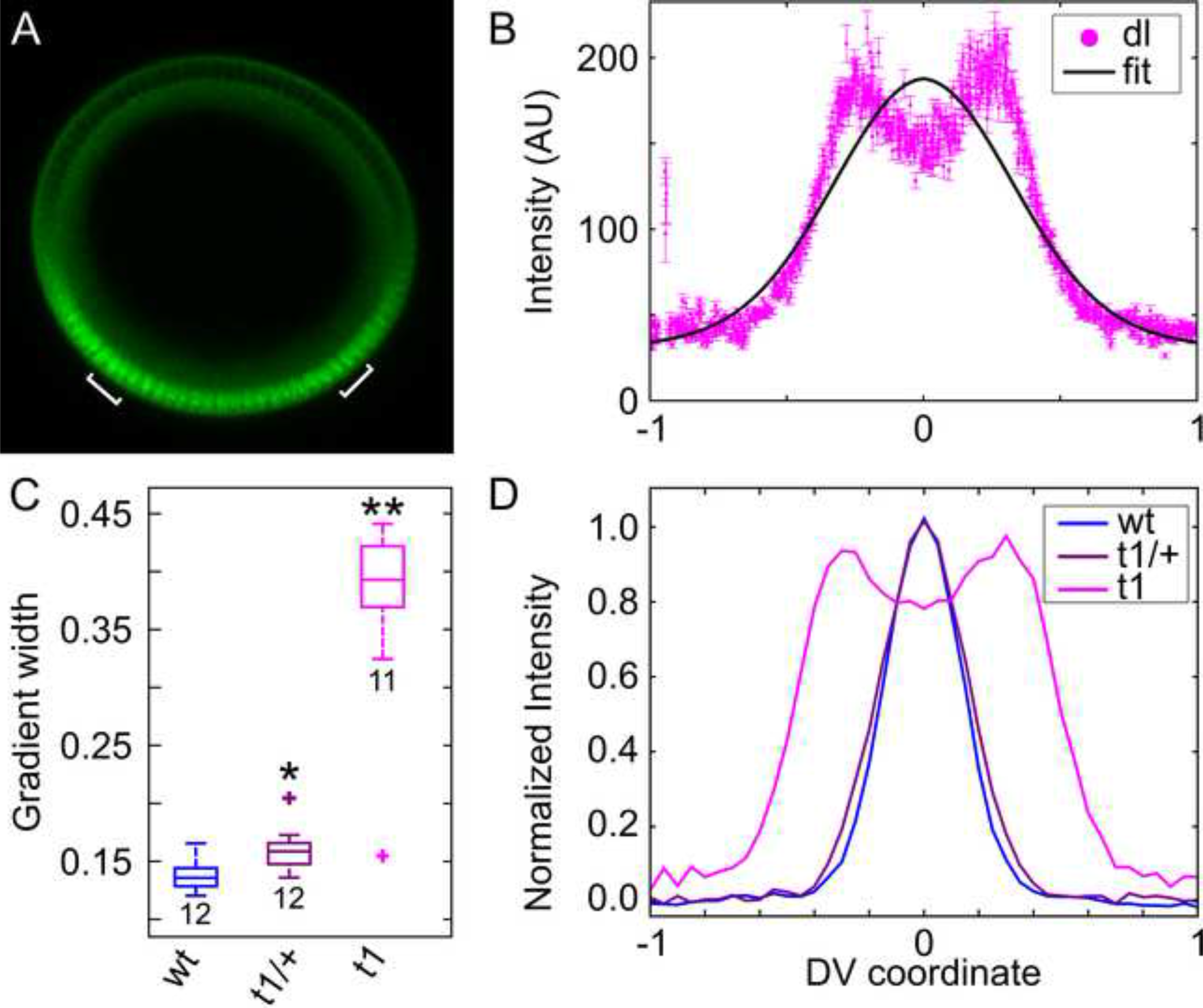
The Hypomorphic Allele *egfr^t1^* Significantly Widens the dl Gradient. (A) Cross-section of an embryo from a mother homozygous for *egfr^t1^* stained for dl. Brackets indicate peaks of nuclear dl. (B) Plotted nuclear intensity of embryo in (A) as a function of DV coordinate (each pink dot is one nucleus). The shape has changed significantly from wild type, as the Gaussian curve (black) does not represent the gradient well. Error bars indicate SEM of nuclear intensity. (C) Box plot of gradient widths (σ) for embryos from mothers with 0 (wt), 1 (t1/+), and 2 (t1) copies of *egfr^t1^.* Numbers indicate sample size. Asterisks indicate statistical significance (* p<0.01, **p<10^−8^) from wt. (D) Normalized average plot of dl gradient in embryos from (C).

### An anteroposterior gradient of dl supports the shuttling hypothesis

It has been suggested that a shuttling phenomenon also occurs through the processing of Spätzle (Spz), the ligand for Toll signaling (Haskel-Ittah et al., 2012), which may explain some of the phenotypes described here. To determine whether the hallmark phenotypes of shuttling could occur without assistance from the protease cascade or Spz processing, we expressed a constitutively active form of Toll (Toll^10b^) at the anterior pole of the developing embryo using the *bicoid* 3′ UTR and the *bicoid* promoter (Huang et al., 1997). Embryos from mothers carrying this construct *(bcd> toll^10b^: bcd 3’ UTR)* have an anterior-posterior (AP) dl gradient in addition to the native DV gradient. Naïvely, one may expect that the existence of two gradients in active Toll signaling would result in higher concentrations of nuclear dl where these two gradients overlap. In contrast, the shuttling hypothesis predicts that dl nuclear concentration would be *lower* in the region of overlap, as the two competing dl/Cact sinks cause Cact to shuttle dl toward both the anterior pole and the ventral midline (see Figure 6A–C). This prediction is borne out in experiment, as these embryos show a decreased intensity of the dl gradient in the region of overlap. Furthermore, 64% (9/14) of these embryos show a visible narrowing of the *sna* expression domain at roughly 30% egg length, consistent with previously published numbers (Huang et al., 1997). There is a dip in dl nuclear intensity at this location as well (Figure 6B,C).

**Figure 6:**
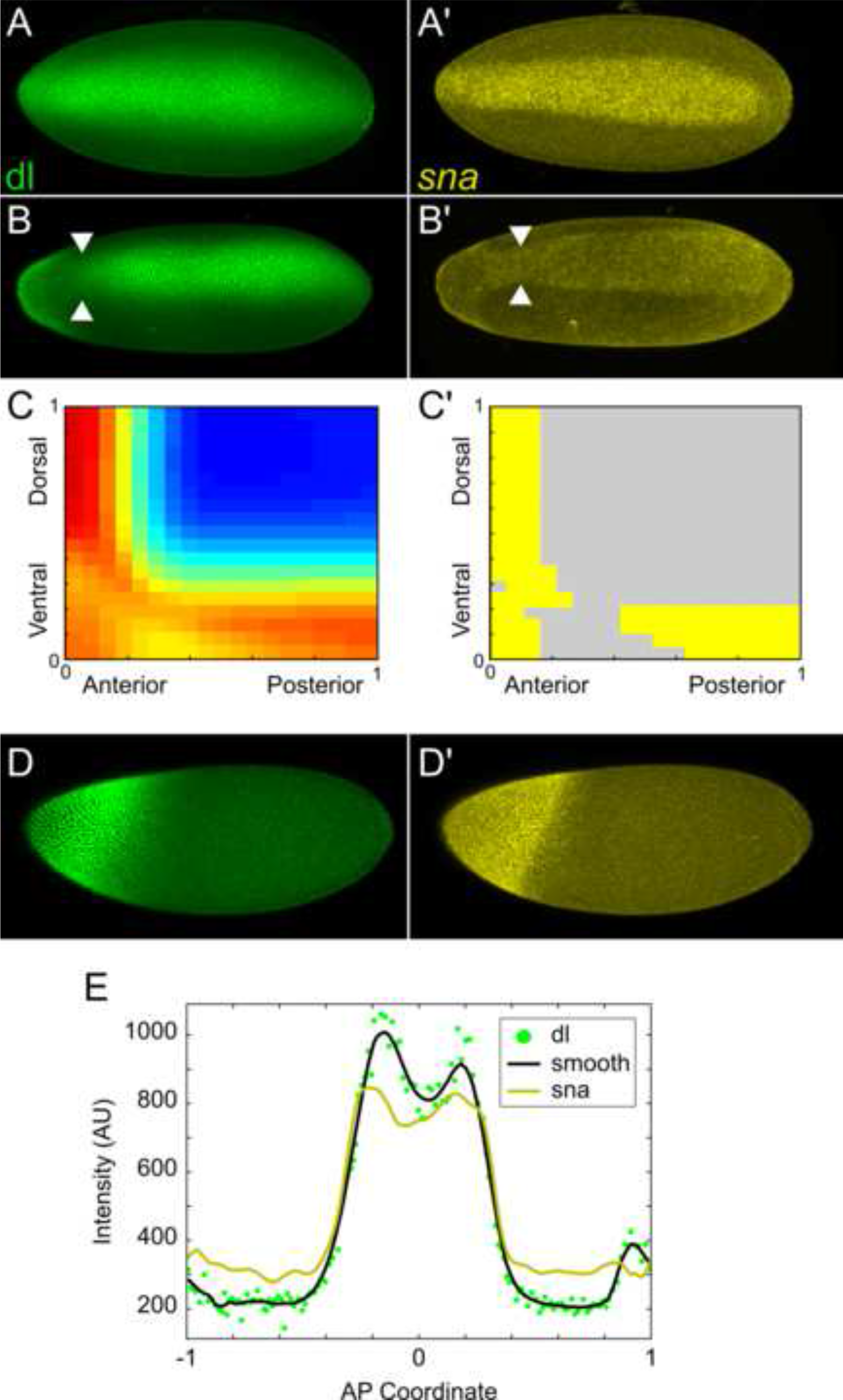
An Ectopic, Anterior-Posterior Dorsal Gradient Exhibits Shuttling Phenomena. (A) and (A′) dl and *sna* expression in a wild type embryo. (B) and (B′) dl and *sna* expression in an embryo with an anteroposterior gradient of dl driven by the *bcd* promoter in addition to the wild type DV gradient. White arrowheads indicate a narrowing of each domain at ~30% EL. (C) and (C′) A 2-D version of the model featuring AP and DV active Toll domains exhibits a competing sink phenotype, where a low point in the dl gradient leads to a gap in *sna* expression. (D) and (D′) dl and *sna* expression in an embryo with an anteroposterior gradient of dl driven by the *hsp83* promoter and the DV gradient abolished by a homozygous mutation in *gd*. (E) Plot of dl and *sna* domains from the embryo in (D). Each green dots is one nucleus and the black curve is a smoothing of the dl data. Embryo images are maximal intensity projections. See also Figure S3.

Our modeling results further support that the shuttling mechanism is responsible for this phenomenon (Figure 6C). We expanded our model system from a one-dimensional array of nuclear compartments to a two-dimensional array and added a second Toll signaling domain perpendicular to the first. Our simulation results show that the overlap between the AP and DV Toll domains creates a low point in nuclear dl concentration, similar to our experimental results. We then approximated a threshold for *sna* expression based on the DV dl gradient, and found the local minimum in dl concentration results in reduced or abolished *sna* expression in that region (Figure 6C′).

We also examined embryos from mothers carrying a homozygous mutation in *gastrulation defective (gd^7^)*, which eliminates the endogenous ventral-to-dorsal gradient. Swapping the *bicoid* promoter for the stronger *hsp83* promoter *(hsp83> toll^10b^: bcd 3’ UTR* construct) (Huang et al., 1997), we were able create a wider dl gradient at the anterior pole. Half of these embryos (7/16) show two peaks of dl both by eye and when the nuclear intensity is plotted (Figure 6D,E). Furthermore, these embryos have a double peak of *sna* (Trisnadi et al., 2013). In order to determine whether or not this double peak phenomenon was a result of the embryo′s geometry at the pole, we analyzed embryos with an AP dl gradient initiated by the weaker *bicoid* promoter. These embryos showed no such double peak effect (see Figure S3). These results further support our shuttling hypothesis as the dl gradient progresses from narrow (weak promoter) to double peak (strong promoter) much as it does in the native DV system when the Toll domain expands.

## Discussion

In this study, we investigated the formation of the dl morphogen gradient in the early *Drosophila* embryo. Based on our model and experimental verification, we conclude that the dl gradient is established by a facilitated diffusion, or “shuttling” mechanism, in which dl/Cact complex diffuses towards the ventral midline where Cact is degraded. Furthermore, our hypothesized mechanism makes sense of several previously unexplained phenomena in the literature.

Besides dl binding to Cact, the shuttling mechanism requires two phenomena to occur. First, dl/Cact complex must be able to move throughout the embryo. Our experiments with dl-paGFP show that dl does indeed move through the embryo in a manner that appears to be consistent with diffusion. In contrast, a previous report showed that each nucleus has its own well-mixed cytoplasmic pool of dl to draw from, and that dl does not readily diffuse from pool to pool (Delotto et al., 2007). This observation was confirmed by a study which showed that barriers to diffusion exist in the early embryo, despite the lack of cell membranes (Mavrakis et al., 2009). However, these results do not contradict our hypothesized mechanism, which relies on movement of dl from one pool to another, because the time scales are different (seconds vs. minutes). Furthermore, observational evidence supports a diffusion-based shuttling mechanism, as lowering the diffusivity of either dl or Cact widens the gradient (Figure 7A), rather than the narrowing that one might expect from a morphogen gradient established by (nonfacilitated) diffusion. The shuttling mechanism explains why dl tagged with a weakly dimerizing GFP widens the gradient more than one tagged with monomeric Venus (Reeves et al., 2012), and also explains the related observation that one copy of dl-mVenus complements loss of endogenous dl while one copy of dl-dGFP does not (Liberman et al., 2009; Reeves et al., 2012). Similarly, this mechanism makes sense of the observation that dl tagged with β-galactosidase, which forms tetramers, is anti- morphic (Govind et al., 1992), as the dl moieties in tetramers of dl-βgal dimerize with endogenous dl to disrupt the formation of the endogenous dl gradient.

**Figure 7:**
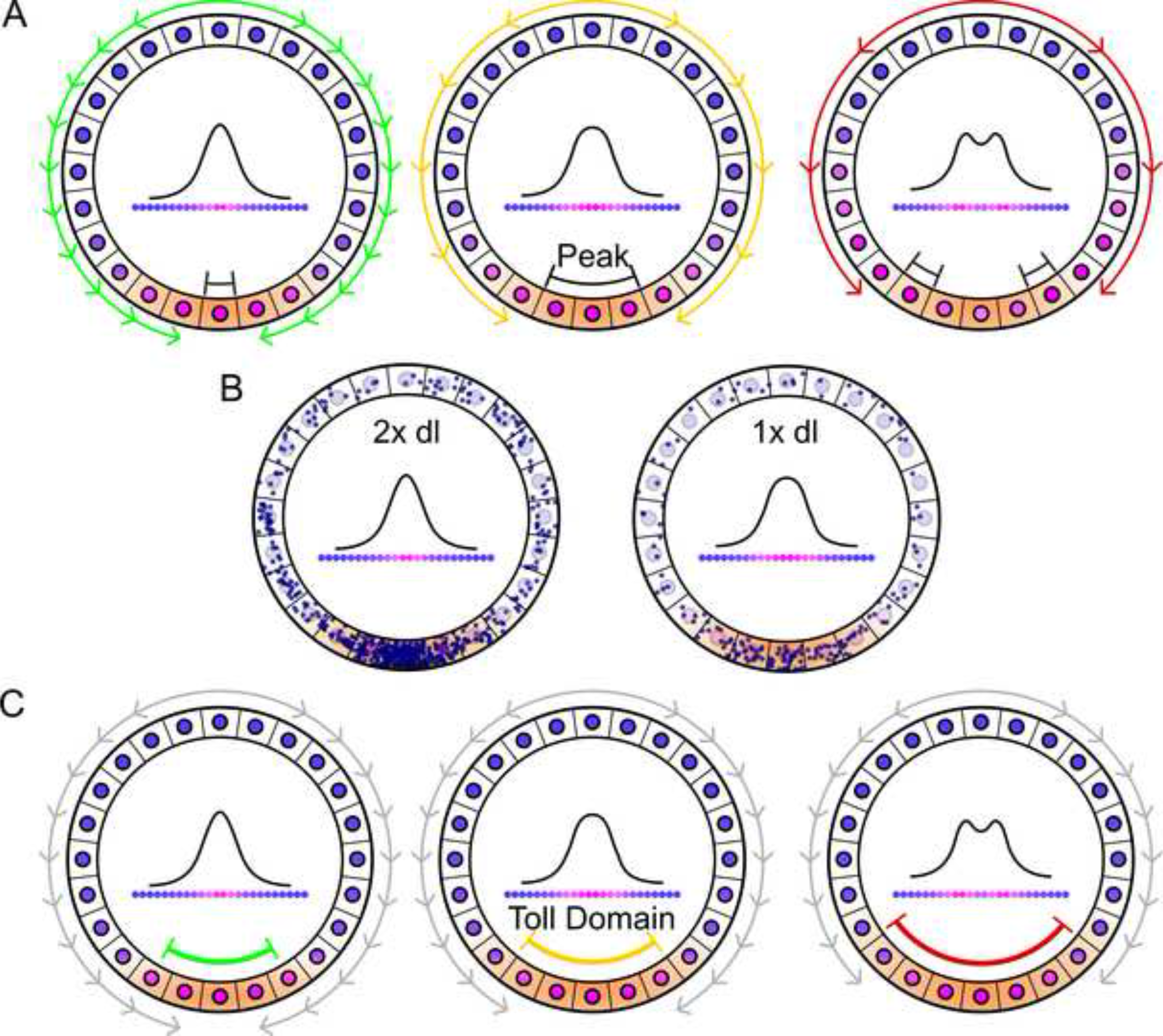
The Shuttling Mechanism Explains the Flat and Double Peak Phenomena, Illustrated in Three Ways: (A) slowing dl/Cact diffusion (colored arrows, from green to red), (B) halving the dosage of dl (blue dots), and (C) expanding the Toll domain (from green to red). Cartoon representative of embryo cross section. Arrows indicate diffusion of dl/Cact complex, Toll domain highlighted in orange, nuclear concentration indicated by blue-pink gradient.

Second, the shuttling mechanism requires the ventral midline to be a sink for the dl/Cact complex. This becomes especially clear in embryos that have an ectopic AP dl gradient in addition to the endogenous DV gradient (Huang et al., 1997). In these embryos, a second sink for the dl/Cact complex is established, and the two competing sinks form a local minimum in the dl gradient where the two overlap, rather than the global maximum one might expect if the two gradients were additive as they would be in a model without facilitated diffusion (Figure 6B,C). This result has been previously observed as a ‘gap’ in the *sna* domain where the two gradients overlap (Huang et al., 1997), similar to our 2D simulation results. These results argue that the Toll domain does act as a sink, a necessary condition for shuttling.

In this system, it appears that active Toll receptors are somewhat limiting as they are easily saturated with dl/Cact complex. This saturation is not essential to the shuttling mechanism *perse*, but it is necessary for the mechanism to explain several observations of the dl/Cact system. Under wild type conditions, a significant flux of dl/Cact complex can bypass the saturated active Toll receptors in the tails of the gradient, which results in the accumulation of a smooth, intense peak of dl signaling at the ventral midline of the embryo. However, in embryos from *dl/+* mothers, there is not enough dl/Cact complex to saturate active Toll receptors in the tails of the gradient, and thus there is less accumulation of dl at the ventral midline (Liberman et al., 2009) (see Figure 7B). Similarly, when the Toll gradient is greatly expanded, as in embryos from *egfr*^11^ mothers (Roth and Schüpbach, 1994), the domains of saturated Toll receptors move further from the ventral midline, resulting in a split peak (Figure 7C). Additionally, the split-peak phenomenon has also been observed in the dl gradient in abnormally large embryos (Garcia et al., 2013). This specific phenotype could be explained by the shuttling of the Toll ligand Spz (Haskel-Ittah et al., 2012); however, we observed the same phenomenon in embryos with an ectopic AP dl gradient established by constitutively active Toll. In both cases, ventrally- (anteriorly-) diffusing dl/Cact complex does not make it to the ventral midline (anterior pole) before being dissociated, leaving the ventral- (anterior-) most nuclei somewhat devoid of dl. A similar mechanism, in which the removal rate of BMP ligands surpasses the rate of BMP flux to the dorsal midline, has been suggested to explain the computationally-predicted split-peak phenotype for the BMP system in the early embryo (Umulis et al., 2006).

The shuttling mechanism has been proposed in several other contexts and in multiple organisms (Ben-zvi et al., 2008; Eldar et al., 2002; Haskel-Ittah et al., 2012; Marqués et al., 1997; Mizutani et al., 2005; Shimmi et al., 2005). For example, during the same stage of *Drosophila* development, shuttling of the BMP ligands Dpp and Scw through the action of Sog and Tsg is responsible for the concentration of BMP signaling to a narrow, intense stripe at the dorsal midline of the embryo. A similar mechanism is also present in BMP signaling in vertebrate embryos to specify the DV axis. In light of these examples, we speculate that this may be a more prevalent mechanism of morphogen gradient formation than previously appreciated.

## Experimental Procedures

### Fly lines

*yw* flies were used as wild type. *dl-paGFP, dl-mGFP,* and *dl-dVenus* were created by BAC recombineering. For live imaging, flies carrying *dl-paGFP* were crossed to flies carrying H2A-RFP on the second chromosome (BS# 23651). *dl/+* flies were created by cleaning up *dl^1^* via two homologous recombinations with yw to generate *dl^1.2.5^. cact/+; cact-full lacZ 25* flies were obtained from David Stein (Fernandez et al., 2001). *dl-lacZ* flies were obtained from Shubha Govind (Govind et al., 1992). *dl-dGFP* and *dl-mVenus* flies and the original BACs used to create them were obtained from Angela Stathopolous. *dl-mGFP,dl^1.2.5^* flies were created by homologous recombination. Presence of *dl-mGFP* was confirmed by w+. Presence of *dl^1.2.5^* was confirmed via sequencing by GENEWIZ, Research Triangle Park, NC. *egfr^t1^/CyO* flies were obtained from the Bloomington Stock Center (# 2079). Flies carrying the *bcd>toll^10b^: bcd 3′UTR* construct were obtained from Angela Stathopolous. The plasmid carrying *FRT-stop-FRThsp83> toll^10b^: bcd 3′UTR* was also obtained from Angela Stathopoulos. To remove the FRT-stop-FRT cassette, we crossed flies carrying this construct into a line carrying hsFLP on the X chromosome (BS# 8862). To remove the native DV dl gradient, flies carrying the *toll^10b^: bcd 3′UTR* construct were crossed into a *gd^7^* background. See Supplemental Experimental Procedures for more details.

### BAC Recombineering

We followed Protocol 3 of the NCI at Frederick Recombineering website
(http://ncifrederick.cancer.gov/research/brb/recombineeringinformation.aspx) to generate dl-paGFP, dl-mGFP, and dl-dVenus in pACMAN (Venken et al., 2006). NEB’s proofreading Q5 DNA polymerase was used to amplify sequences at a high level of authenticity. Using a GalK selection protocol (Warming et al., 2005), single amino acid mutations were introduced at residue 206 in previously established BACs (Reeves et al., 2012) to generate dl-mGFP (A206K) and dl-dVenus (K206A). dl-paGFP was created by adding the open reading frame of *paGFP* (Addgene plasmid #11911),(Patterson and Lippincott-Schwartz, 2002) in frame to the 3′ end of the *dl* open reading frame in a dl rescue construct (Reeves et al., 2012) in pACMAN (Venken et al., 2006) using a 6x-Gly linker. See Table S1 for a list of primers used.

### *Fluorescent* in situ *Hybridization*

All embryos were aged to 2-4 hours, except for “young” yw embryos, which were aged to 0-2 hours, then fixed in 37% formaldehyde according to standard protocols (Kosman et al., 2004). A combination fluorescent *in situ* hybridization/fluorescent immnuostaining was performed according to standard protocols (Kosman et al., 2004). Briefly, fixed embryos were washed in Tween/PBS buffer, then hybridized with anti-sense probes at 55 ^°^C overnight. The embryos were then washed and incubated with primary antibodies at 4 ^°^C overnight. Next, the embryos were washed and incubated for 1-2 hrs with fluorescent secondary antibodies at room temperature. The embryos were then washed and stored in 70% glycerol at −20 ^°^C. Embryos were imaged within one month of completing the protocol. See Supplemental Experimental Procedures for more details.

### Mounting and Imaging of Fixed Samples

Embryos were cross sectioned and mounted in 70% glycerol as described previously (Carrell and Reeves, 2015). Briefly, a razor blade was used to remove the anterior and posterior thirds of the embryo, leaving a cross section roughly 200 μm long by 200 μm in diameter. These sections were then oriented such that the cut sides became the top and bottom. They were then imaged at 20x on a Zeiss LSM 710 microscope. 15 z-slices 1.5 μm apart were analyzed. Embryos with an AP Dorsal gradient were mounted laterally in 70% glycerol using one piece of double-sided tape (weak *bicoid* promoter) or two pieces of double-sided tape (strong *hsp83* promoter). Images were taken at 2.5 μm intervals from just above the top of the embryo to the depth at which the embryo reached maximal size in the xy plane, which was assumed to be the midsagittal section. Stacks ranged from 15-25 slices.

### Data Analysis

Images of embryo cross sections were analyzed using previously derived code (Trisnadi et al., 2013). Briefly, the border of the embryo was found computationally, then the nuclei were segmented using a local thresholding protocol. The intensity of dl in each segmented nucleus was calculated as the ratio between the intensity in the dl channel divided by the intensity in the nuclear channel. The intensity of mRNA expression was calculated as average intensity within an annulus roughly 18 μm wide around the perimeter of the embryo. A description of the image analysis of whole mount embryos can be found in the Supplemental Experimental Procedures.

All dl gradients were fit to a Gaussian, and these fits were used to determine the width parameter, σ. Normalized intensity plots were generated by fitting each embryo’s data to its own Gaussian by subtracting the B value and 70% of the M value, then dividing by the A value. (X = (x − B − 0.7M)/A)). The average normalized intensity plot was generated by averaging the plots of all embryos in the specified genotype.

Statistical significance was calculated using two-tailed homoscedastic t-tests.

### Activating paGFP in Live Embryos

Embryos were dechorionated by hand or for 30 s in 100% bleach. They were then mounted laterally on a slide coated with heptane glue. Deionized water was used as a mounting medium. Two pieces of double-sided tape were used to attach the coverslip. Images were taken using a 40x water immersion objective on an LSM 710 confocal microscope. Activation box: 9000 pixels (300 microns × 30 microns), number of activation passes: 10, rest time: 15 s, number of cycles: 40, laser power: 3% for embryo activated near ventral midline, 2.5% for embryo activated near dorsal midline. A 405 nm laser was used for activation and a 488 nm laser was used for excitation of the activated GFP.

### Model of dl/Cact interactions

We constructed a model of dl/Cact interactions consisting of eight differential equations, representing nuclear and cytoplasmic dl, Cact and dl/Cact, as well as activated Toll receptors and Toll receptors bound to dl/Cact complex (Toll:dl/Cact). The equations were then non-dimensionalized, resulting in a model with 20 free parameters. Interactions were simulated during nuclear cycles 10-14, as previously published (O′Connell and Reeves, 2015). Optimization was performed in MATLAB^®^ using an evolutionary optimization algorithm with stochastic ranking (Runarsson and Yao, 2000, 2005), yielding 114 parameter sets that were generally consistent with the spatiotemporal data published by Reeves et al. (2012). Further detail on model equations can be found in the Supplementary Experimental Procedures.

Adding together the model equations for dl and dl/Cact (Eqns 1-4 in Supplemental Experimental Procedures), we obtain the following equation for total dl:

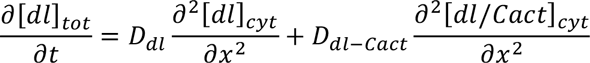

where *D_i_* represents the diffusion coefficient for species *i*.

## Author Contributions

SNC, AEA, and SMS conducted experiments. MDO conducted mathematical modeling. SNC, MDO, and GTR wrote the manuscript. SNC and MDO generated figures.

## Acknowledgments

Thanks to Angela Stathopoulos and Leslie Dunipace for provision of dl-dGFP and dl-mVenus flies and BACs, *bcd> toll^10b^: bcd 3’ UTR* flies, and the DNA used to generate *hsp83> toll^10b^: bcd 3’* UTRflies. Thanks to ImmunoReagents for providing their goat ant-biotin antibody. We are greatful to Alex Thomas for generating the sna-biotin probe. Thanks to Parv Gondalia for generating the sna-fitc probe. Thanks to David Stein for providing the cact-lacZ fly line. We are grateful to Shubba Govind for providing the dl-lacZ flies. During this work, SNC and GTR (partially) were supported by a CAREER award funded by the United States National Science Foundation (www.nsf.gov), grant number CBET-1254344; MDO was supported by a Graduate Assistance in Areas of National Need Fellowship in Computational Science from the United States Department of Education (www.ed.gov), grant number P200A120047.

